# Genomic and physiological characterization of *Neobacillus ammoniipjilus* sp. nov. reveals versatile inorganic nitrogen utilization and ammonia-mitigation potential

**DOI:** 10.64898/2026.10.12.758474

**Authors:** Thi Tuyet Nhan Le, K.C Rabin, Seung-Woo Jeong, Jaisoo Kim

**Author notes:** Thi Tuyet Nhan Le and Rabin K.C. contributed equally to this work. Corresponding author: Jaisoo Kim.

## Abstract

A novel bacterial strain (KR4-4ᵀ), was isolated from soil in the Republic of Korea and subjected to polyphasic taxonomic and functional characterization. Phylogenetic analysis based on the 16S rRNA gene sequence placed strain KR4-4ᵀ within the genus *Neobacillus*. Phylogenomic analysis based on 92 bacterial core genes further supported its distinct taxonomic position. Average nucleotide identity and digital DNA–DNA hybridization values between KR4-4ᵀ and related *Neobacillus* type strains were 69.76–82.31% and 18.3–25.9%, respectively, supporting its recognition as a distinct species. Cells were Gram-stain-positive, non-motile, facultatively anaerobic rods, with growth occurring at 15–45 °C, pH 6.0–8.0, and 0–2.0% (w/v) NaCl. The major cellular fatty acids were iso-C_15:0_ and anteiso-C_15:0_, while the major polar lipids included diphosphatidylglycerol, phosphatidylglycerol, and phosphatidylethanolamine. Genome analysis revealed genes associated with inorganic nitrogen transport, nitrate and nitrite reduction, and ammonium assimilation. In liquid culture, KR4-4ᵀ removed NH₄⁺-N under aerobic conditions and was able to utilize ammonium, nitrate, and nitrite as individual nitrogen sources, with removal efficiencies of 86.5%, 69.35%, and 62.9%, respectively. The strain also contributed to the reduction of gaseous NH₃ in a closed-flask assay. Together, these physiological and genomic characteristics indicate a versatile capacity for inorganic nitrogen utilization and transformation and suggest potential for further evaluation in biological ammonia-mitigation applications. Based on phylogenetic, genomic, phenotypic, and chemotaxonomic evidence, strain KR4-4ᵀ represents a novel species of the genus *Neobacillus*, for which the name *Neobacillus ammoniiphilus* sp. nov. is proposed. The type strain is KR4-4ᵀ (= KACC 24003ᵀ = CCTCC AB 2025077ᵀ).

**Importance:** *Neobacillus* species occur in diverse environments, but their capacity to use and transform multiple inorganic nitrogen forms remains poorly characterized. This study describes *Neobacillus ammoniiphilus* sp. nov. strain KR4-4ᵀ and links its taxonomic and genomic features with physiological nitrogen-utilization phenotypes. The strain used ammonium, nitrate, and nitrite as individual nitrogen sources, showed substantial ammonium removal under aerobic conditions, and enhanced the reduction of gas-phase ammonia in a sealed-flask assay. Genome analysis identified systems associated with inorganic nitrogen transport, nitrate and nitrite reduction, ammonium assimilation, and nitrogen regulation. These findings expand knowledge of nitrogen metabolism within *Neobacillus* and identify KR4-4ᵀ as a candidate for further evaluation in biological ammonia-mitigation applications.

## Introduction

As a critical building block for all living organisms, nitrogen acts as a key determinant of biological productivity across diverse ecosystems. However, anthropogenic activities have led to excessive inputs of reactive nitrogen compounds into terrestrial and aquatic ecosystems, severely disrupting the natural nitrogen cycle. Specifically, the unregulated discharge of ammonium (NH_4_^+^), nitrate (NO_3_^-^), and nitrite (NO_2_^-^) from agricultural, industrial, and domestic effluents accelerates eutrophication and severely degrades water quality, underscoring the critical need for advanced nitrogen management and remediation strategies (1, 2). Concurrently, atmospheric ammonia (NH_3_) emissions, predominantly derived from agricultural and livestock sectors, pose severe environmental risks by driving ecosystem acidification and facilitating the formation of secondary fine particulate matter (PM2.5) (3). Microbial communities play a pivotal role in mediating the interconnected pathways of the nitrogen cycle, including ammonium assimilation, nitrification, denitrification, and dissimilatory nitrate reduction to ammonium (1, 4). Metabolically versatile heterotrophic microorganisms that can perform simultaneous nitrogen transformations under fully aerobic conditions have garnered significant attention. These strains offer a streamlined approach to biological nitrogen removal by consolidating multi-stage processes into a single reactor while concurrently promoting rapid biomass production (5–7). Despite these advantages, metabolic pathways in aerobic heterotrophs are highly complex, and phenotypic observations alone often fail to fully elucidate the underlying biochemical mechanisms. Recent studies have identified an increasing number of aerobic heterotrophic bacteria capable of utilizing multiple inorganic nitrogen sources via coupled pathways involving assimilation, heterotrophic nitrification, and aerobic denitrification (8–10). For instance, *Halomonas venusta* SND-01 demonstrated efficient simultaneous removal of ammonium, nitrate, and nitrite, with nitrogen-balance and genomic analyses confirming the co-activation of ammonium assimilation and heterotrophic nitrification–aerobic denitrification (9). Similarly, *Klebsiella* sp. TSH15 was reported to clear ammonium and nitrate through integrated pathways of assimilatory/dissimilatory nitrate reduction and ammonium assimilation, validated by whole-genome sequencing and transcriptional profiling of nitrogen-metabolism genes (10). These findings highlight that nitrogen mitigation by individual bacterial isolates often relies on a network of interconnected metabolic routes rather than a single canonical pathway. Nagel and Andreesen first proposed the genus *Neobacillus* in 1991 as a member of the genus *Bacillus* (11). Patel and Gupta (2020) reclassified it as the type species of the genus *Neobacillus* a member of the family *Bacillaceae* after conducting comprehensive phylogenomic and comparative genomic analyses to address the complex, multifaceted nature of the vast *Bacillus* genus (12). At the time of writing, twenty-nine species have been validly published in this genus (https://lpsn.dsmz.de/genus/neobacillus). Species within *Neobacillus* are ubiquitous in diverse habitats and exhibit profound physiological and metabolic plasticity (13–16). Notably, *Neobacillus nitrireducens* OS1-33 was recently described as a novel soil bacterium capable of reducing nitrate under low-temperature, microaerobic conditions, confirming nitrogen-transforming traits within the genus (17). Nevertheless, the capacity of *Neobacillus* strains to concurrently utilize and transform diverse inorganic nitrogen species-particularly under strictly aerobic conditions-remains poorly understood. Consequently, integrating taxonomic, physiological, and genomic data is essential to map the functional landscape of nitrogen metabolism within this genus. In the present study, strain KR4-4^T^ was characterized using a polyphasic taxonomic approach encompassing phylogenetic, phylogenomic, phenotypic, and chemotaxonomic analyses. The kinetics of ammonium, nitrate, and nitrite utilization by strain KR4-4^T^ were systematically evaluated under aerobic conditions, tracking temporal dynamics in nitrogen profiles, cell proliferation, and biomass yield. Furthermore, its potential to mitigate gaseous ammonia was assessed, complemented by whole-genome annotation to identify functional genes governing nitrogen metabolism and environmental adaptation. The primary objectives of this study were to resolve the precise taxonomic position of strain KR4-4^T^ and to elucidate its inorganic nitrogen-utilization mechanisms through the lens of its genomic potential. Based on polyphasic evidence, strain KR4-4^T^ represents a novel species within the genus *Neobacillus*, for which the name *Neobacillus ammoniiphilus* sp. nov. is proposed.

## Results and Discussion

### Identifying the strain

The 16S rRNA gene sequence of strain KR4-4^T^ was 1,564 bp in length. The closest match for the 16S rRNA gene sequence was *Neobacillus drentensis* LMG 21831^T^ with a similarity of 97.8%, which is below the species delineation threshold of 98.7% (18). Phylogenetic analyses of 16S rRNA gene sequences also showed that strain KR4-4^T^ formed a stable branch with *Neobacillus drentensis* LMG 21831^T^ in NJ, ML, and ME (Figures S1, S2, and S3). A comparative genomic analysis, including the construction of a phylogenetic tree based on 92 bacterial core genes, was conducted to further clarify the taxonomic position of strain KR4-4^T^ (Figure 1). Core genome-based phylogeny revealed that strain KR4-4^T^ formed a tight cluster with *Neobacillus drentensis* LMG 21831^T^ and a large group with *Neobacillus soli* DSM 15604^T^. These strains formed a tight cluster, and they were selected as reference species for further comparative analysis.

**Figure 1.**
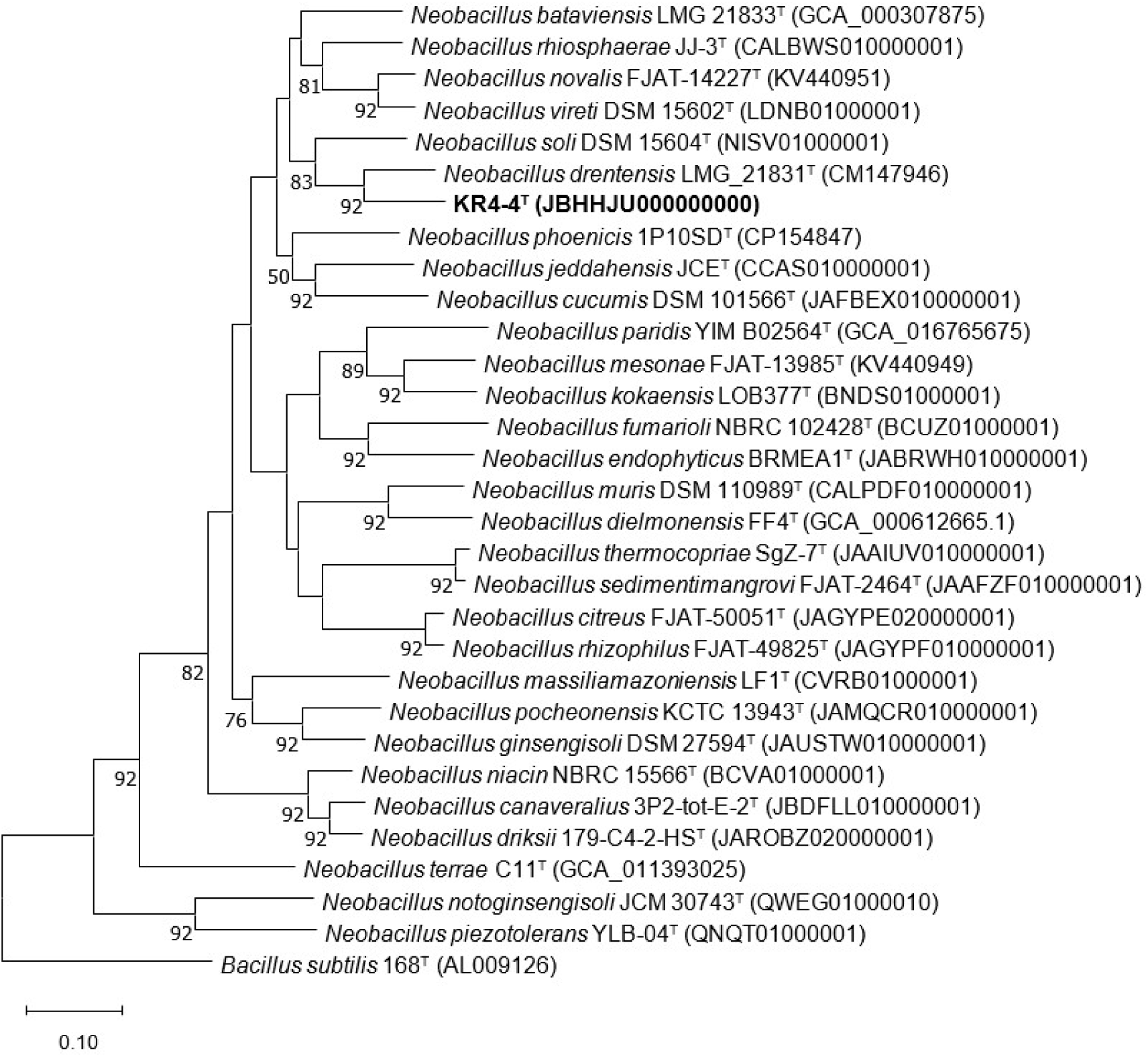
The core genome-based phylogenetic tree was reconstructed by the UBCG phylogenomics pipeline (concatenated alignment of 92 core genes) for KR4-4^T^ and members of the genus *Neobacillus*. *Bacillus subtilis* was used as an outgroup. GenBank accession numbers are given in parentheses. Scale bar, 0.10 nucleotide substitutions per site.

The ANI values between strain KR4-4^T^ and the type strains in the genus *Neobacillus* ranged from 69.76 to 82.31 %, while the dDDH values ranged from 18.3 to 25.9 % (Table S1). Among them, *Neobacillus drentensis* LMG 21831^T^ had the highest ANI and dDDH values with KR4-4^T^. This further confirmed and strengthened their close relationship, as shown by the phylogenetic tree based on 16S and the core genome. These values were well below the commonly accepted species delineation thresholds, which were 95–96% (19) for ANI and 70% for dDDH (20). These values indicate that KR4-4^T^ is separated from all known taxa at the species level and represents a novel *Neobacillus* species.

### Physiology and chemotaxonomic characterization

Strain KR4-4^T^ was found to be negative for both catalase and oxidase, non-motile, and a facultative anaerobic bacterium. Cells were rod-shaped (0.5-1.5 × 0.9-4.8 µm) (Figure S4). Growth was detected at temperatures ranging from 15 to 45 °C (optimum 30-40 °C), pH levels from 6.0 to 8.0 (optimum 7.0-7.5), and at NaCl concentrations from 0 to 2.0% (w/v). It demonstrated the ability to hydrolyze starch, DNase, esculin and gelatin, but not casein and urease. Nitrate is reduced to nitrite. In the API ZYM test, strain KR4-4^T^ tested positive for esterase (C4), esterase lipase (C8), leucine arylamidase, α-chymotrypsin, β-galactosidase, β-glucuronidase, and α-glucosidase. The utilisation of the sole carbon source (API 20NE and API 50CH) was positive for D-glucose, D-mannose, D-mannitol, N-acetyl-glucosamine, D-maltose, malate, phenylacetic acid, D-lactose, D-saccharose, D-trehalose, D-melezitose, amidon, and glycogene. The distinct physiological and biochemical characteristics of strain KR4-4^T^ compared with its closest phylogenetic neighbours are depicted in Table 1 and Table S2.

**Table 1.** Differential characteristics of 1, KR4-4T, compared with 2, *Neobacillus drentensis* KACC 12118^T^ and 3, *Neobacillus soli* KACC 12115^T^ (data are from the present study). +, positive; –, negative; +/−, weak positive or ambiguous.

| Characteristics | 1 | 2 | 3 |
| --- | --- | --- | --- |
| Temperature range for growth (°C) | 15-45 | 10-45 | 20-40 |
| Temperature optimum for growth (°C) | 30-40 | 25-35 | 25-30 |
| pH range for growth | 6.0-8.0 | 6.0-8.5 | 5.5-8.5 |
| pH optimum for growth | 7.0-7.5 | 7.0-7.5 | 7.0-7.5 |
| Highest salt tolerance (% w/v) | 2.0 | 3.5 | 3.0 |
| Hydrolysis of: |  |  |  |
| Casein | - | - | + |
| Gelatin | + | - | + |
| Starch | + | - | + |
| Esculin | + | + | - |
| Enzyme activities |  |  |  |
| Esterase lipase (C8) | + | - | + |
| Valine arylamidase | +/- | - | + |
| β-glucosidase | - | + | - |
| Arginine dihydrolase | - | + | - |
| Utilisation of: |  |  |  |
| D-Mannitol | + | - | - |
| Adipic acid | - | - | + |
| Malate | + | - | + |
| Trisodium citrate | - | - | + |
| Phenylacetic acid | + | - | + |
| D-Galactose | - | +/- | + |
| D-mannose | + | - | +/- |
| Salicine | +/- | + | - |
| D-lactose | + | + | - |
| D-Melibiose | - | + | - |
| D-Melezitose | + | + | - |
| D-Raffinose | - | + | - |
| Glycogene | + | - | + |

The major cellular fatty acids (>5%) were iso C_15:0_ (35.3%), anteiso C_15:0_ (33.5%), C_16:0_ (5.9%), and anteiso C_17:0_ (5.1%). The fatty acid profile was similar to that of *Neobacillus drentensis* KACC 12118^T^ and *Neobacillus soli* KACC 12115^T^; however, there were differences in the proportions of primary fatty acids and in the presence or absence of minor fatty acids (Table S3). The major polar lipids of strain KR4-4^T^ include diphosphatidylglycerol (DPG), phosphatidylglycerol (PG), phosphatidylethanolamine (PE), unidentified aminolipid (AL), and an unidentified phospholipid (PL) were also detected in minor amounts (Figure S5).

### Genomic characterization and analysis

The assembled draft genome sequence of strain KR4-4^T^ was 5,632,133 bp in size and contained an N50 of 163,402 bp for the filtered reads, 45 rRNA genes, 151 tRNA genes, and 5,603 genes in total, including 5,334 protein-coding genes. The DNA G+C percent was 38.5 % (Table S4). The Busco analysis showed that completeness of the KR4-4^T^ genome was 100%. Annotations from the KEGG database showed that a total of 2,851 genes were annotated, representing 53.5% of the genome (Figure S6). The functional profiling revealed that a substantial portion of these annotated genes was involved in metabolism. Aside from genes assigned to general metabolic overviews (1,723), the most prominent metabolic subgroups included carbohydrate metabolism (342), amino acid metabolism (297), metabolism of cofactors and vitamins (168), and energy metabolism (158). Additionally, within the genetic information processing category (199 genes), translation (82) and replication and repair (72) constituted the largest functional classes. Other notable functional categories were environmental information processing (238 genes), primarily driven by membrane transport (131), and cellular processes (111 genes). The genes associated with nucleotide metabolism (90) and amino acid metabolism (297) may play vital roles in nitrogen metabolism or specific environmental adaptation in the studied strain.

Genomic data analysis using antiSMASH identified a total of 10 biosynthetic gene clusters (BGCs) in the genome of strain KR4-4^T^ (Table S5). These include various secondary metabolite regions, such as terpenes, T3PKS, and RiPPs. The lack of homologous matches for other predicted BGCs suggests the presence of novel metabolic pathways within this strain. Notably, a high-confidence lassopeptide gene cluster (Region 30.1) showing strong homology to paeninodin, a peptide with potent antimicrobial activity, was detected. This finding suggests that strain KR4-4^T^ is capable of secreting antimicrobial substances for self-defence, potentially contributing to its survival competitiveness and environmental adaptation.

### Genomic insights into nitrogen metabolism and heat stress adaptation

#### Functional genes associated with nitrogen metabolism in strain KR4-4ᵀ

Genome annotation and KEGG pathway reconstruction identified 22 genes associated with nitrogen metabolism in strain KR4-4ᵀ (Figure 2; Table S6). These genes encompass functions related to inorganic nitrogen transport, nitrate and nitrite reduction, ammonium assimilation, nitrogen regulation, and the response to reactive nitrogen species. Genes encoding transport systems for nitrate (NRT2), nitrite (*nirC*), and ammonium (*amt*) were identified, suggesting the capacity of KR4-4ᵀ to acquire multiple inorganic nitrogen sources from the environment.

**Figure 2.**
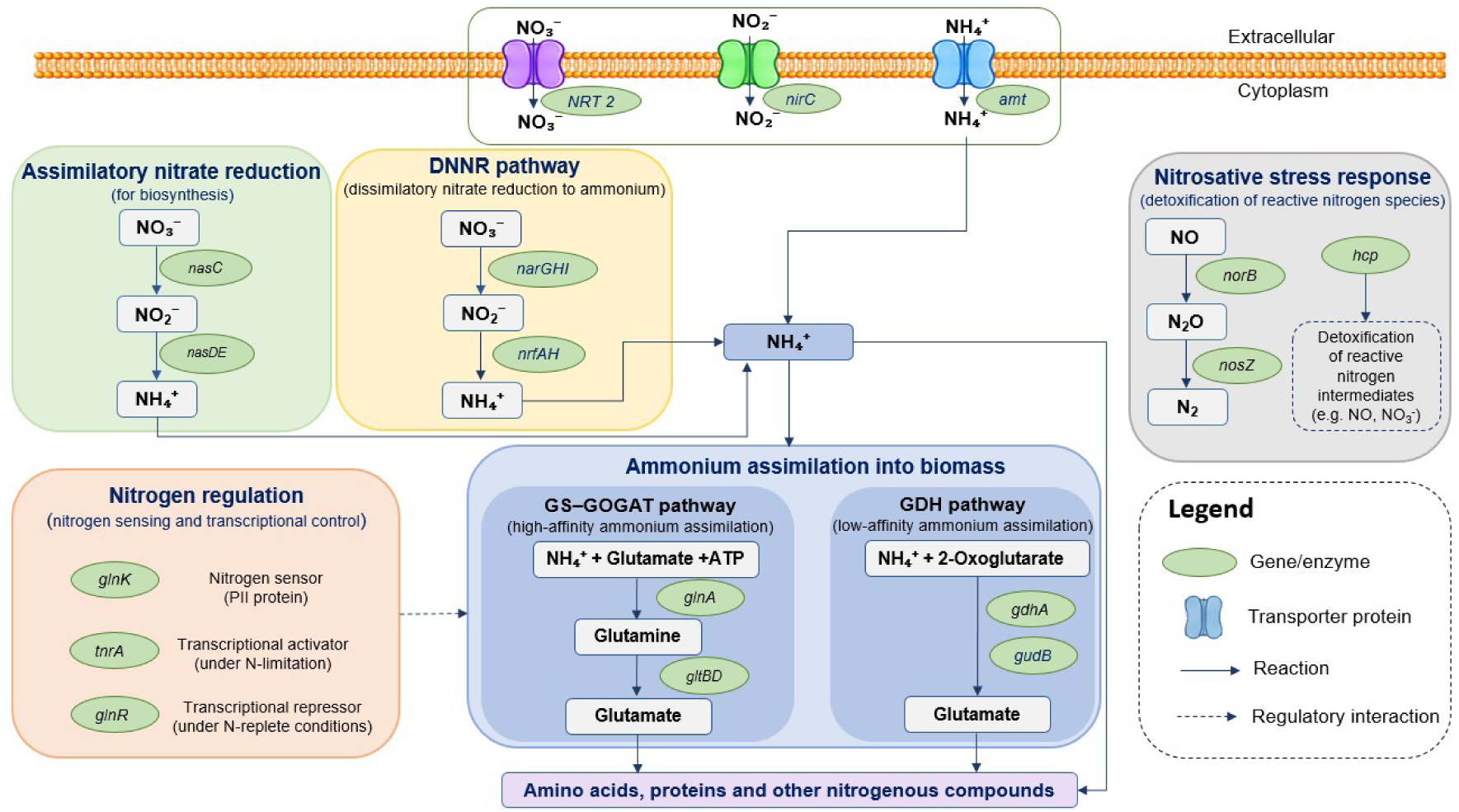
Schematic diagram of the nitrogen metabolism pathway of strain KR4-4^T^.

Genes associated with both assimilatory and dissimilatory nitrate/nitrite reduction were also detected. The assimilatory pathway was represented by *nasC* and *nasDE*, which are involved in the sequential reduction of nitrate to nitrite and subsequently to ammonium for biosynthetic purposes. In parallel, *narGHI* and *nrfAH* were identified, providing genomic evidence for the potential dissimilatory reduction of nitrate and nitrite to ammonium (21). Together, these pathways indicate that oxidized inorganic nitrogen compounds may serve as potential sources of ammonium for subsequent cellular assimilation.

The resulting ammonium can be incorporated into cellular biomass through two major assimilation routes. Genes encoding the high-affinity GS–GOGAT system, including *glnA*, *gltB*, and *gltD*, were identified, together with the glutamate dehydrogenase genes *gdhA* and *gudB*, which provide an alternative route for ammonium assimilation. Consistent with these metabolic functions, the nitrogen regulatory genes *glnK*, *tnrA*, and *glnR* were also detected, indicating genetic capacity for coordinating nitrogen uptake and assimilation in response to nitrogen availability (22–24).

In addition, *hcp*, *norB*, and *nosZ*, which are associated with reactive nitrogen metabolism and/or detoxification, were identified (25). Their presence suggests a potential capacity to cope with nitrosative stress and reactive nitrogen intermediates. However, canonical nitrite reductase genes involved in denitrification, such as *nirK* and *nirS*, were not detected by KEGG. Therefore, the occurrence of *norB* and *nosZ* alone should not be interpreted as evidence for a complete canonical denitrification pathway in KR4-4ᵀ.

Overall, the genomic repertoire of KR4-4ᵀ supports metabolic potential for the uptake and utilization of multiple inorganic nitrogen forms, their conversion to ammonium, and subsequent incorporation into biomass. In particular, the *nasC*, *nasDE, narGHI* and *nrfAH* gene sets provide potential routes linking oxidized nitrogen compounds to ammonium, whereas the GS–GOGAT and GDH systems provide complementary mechanisms for ammonium assimilation into cellular nitrogenous compounds.

#### Genomic features potentially associated with heat-stress responses in strain KR4-4ᵀ

Genome annotation identified several genes potentially associated with protein homeostasis and stress responses in strain KR4-4ᵀ (Table S7). These included the major molecular chaperone systems *dnaK*, *dnaJ*, *grpE*, *groEL* and *groES*, the stress-responsive chaperone gene *hslO*, and genes encoding protein quality-control and proteolytic components, including *clpB*, *clpC*, *clpE*, *clpP*, *clpX*, *lon*, *hslV*, *degP*, and *ftsH*. Genes encoding the stress regulators *hrcA*, *ctsR*, and *sigB* were also detected. These systems are associated with protein folding, removal of damaged proteins, and cellular stress responses in bacteria (26–28), suggesting a genomic capacity of KR4-4ᵀ to maintain protein homeostasis under stressful conditions. Together with the observed growth of KR4-4ᵀ at temperatures up to 45 °C, these genomic features suggest that the strain may tolerate moderately elevated temperatures, a trait that could be advantageous for ammonia-removal applications under temperature-variable or relatively warm conditions. However, ammonia removal by KR4-4ᵀ at elevated temperatures was not evaluated in this study and requires further investigation.

### Ammonium removal and nitrogen transformation by strain KR4-4T

The ammonium-removal capacity of strain KR4-4ᵀ and the associated changes in inorganic nitrogen species were evaluated under aerobic conditions in ANM containing glucose as the carbon source (Fig. 3). In the inoculated treatment, NH₄⁺-N decreased progressively from 184.13 mg/L at 0 h to 159.00 mg/L at 24 h and 104.03 mg/L at 48 h, reaching 98.38 mg/L at 96 h. In contrast, NH₄⁺-N remained comparatively stable in the uninoculated control, decreasing only from 184.32 to 174.08 mg/L. Thus, the inoculated culture showed a net decrease in NH₄⁺-N of 85.75 mg/L after 96 h.

**Figure 3.**
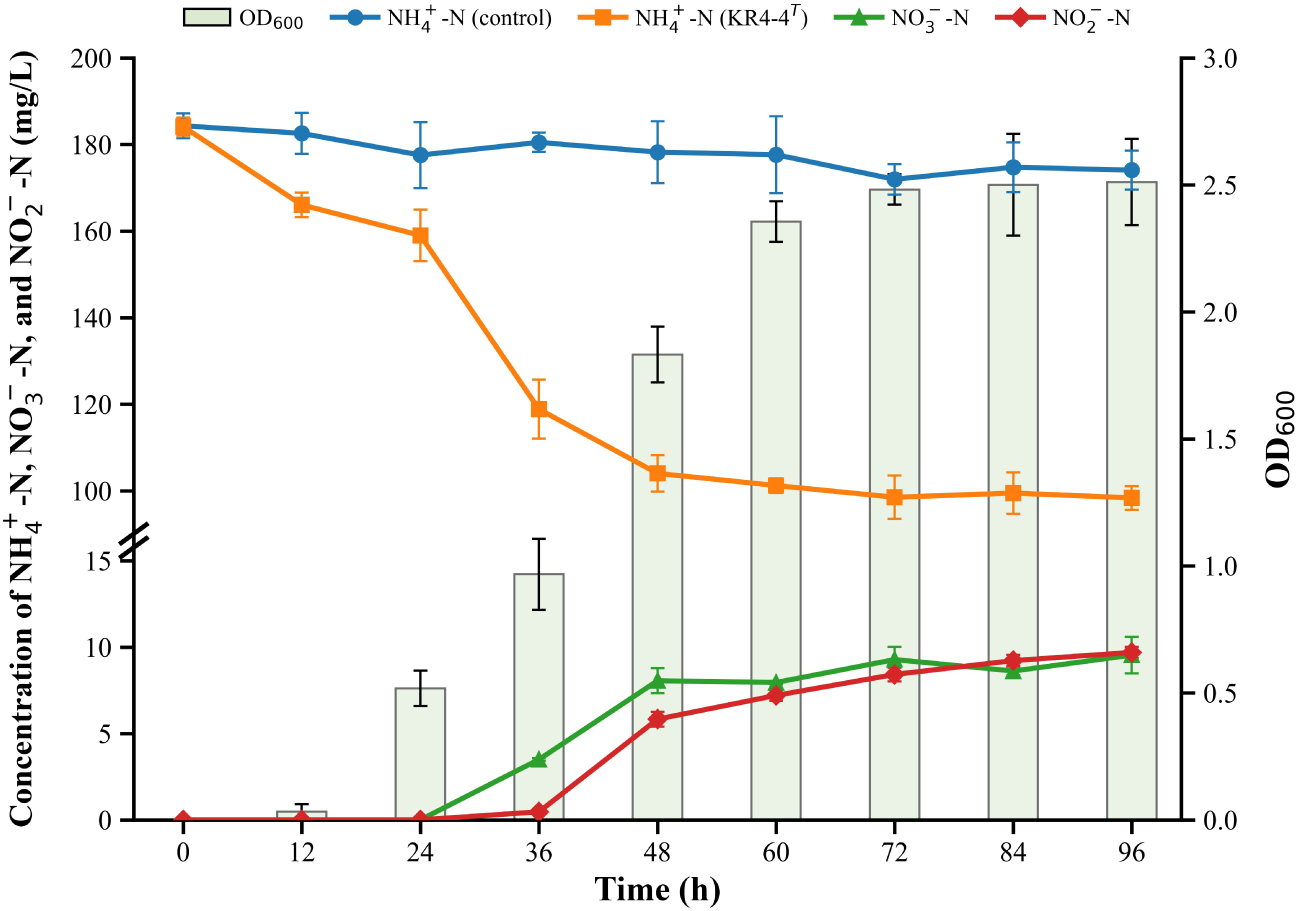
Ammonium removal and nitrogen transformation by strain KR4-4ᵀ. Changes in NH₄⁺-N, NO₂⁻-N, NO₃⁻-N, and OD₆₀₀ were monitored over 96 h. NH₄⁺-N concentrations in uninoculated medium were included as a control. Data are presented as mean ± SD (n = 3).

The NH₄⁺-N decrease was accompanied by substantial bacterial growth. OD₆₀₀ increased from approximately 0 at 0 h to 0.52 at 24 h, 1.83 at 48 h, and 2.51 at 96 h. The most pronounced decrease in NH₄⁺-N occurred between 24 and 48 h, coinciding with rapid cell growth, suggesting that ammonium utilization was closely associated with biomass formation. This interpretation is supported by the genomic identification of the ammonium transporter amt and the GS–GOGAT (*glnA*, *gltB*, and *gltD*) and GDH (*gdhA* and *gudB*) systems, which provide potential routes for ammonium uptake and assimilation into central nitrogen metabolism. Assimilation has similarly been reported as an important route of ammonium removal in aerobic heterotrophic bacteria (9, 29).

Changes in oxidized nitrogen species were also observed during ammonium removal. NO₃⁻-N and NO₂⁻-N were not detected during the first 24 h but became detectable at 36 h and subsequently accumulated to 9.54 and 9.69 mg/L, respectively, at 96 h. Their appearance indicates that ammonium removal was accompanied by nitrogen transformation rather than biomass assimilation alone. However, the accumulated NO₃⁻-N and NO₂⁻-N together accounted for only approximately 19.2 mg/L N, substantially less than the 85.75 mg/L decrease in NH₄⁺-N, suggesting that these oxidized products accounted for only part of the removed ammonium.

Genome analysis further revealed *nasC*, *nasDE*, *narGHI*, and *nrfAH*, indicating genetic potential for assimilatory and dissimilatory nitrate/nitrite reduction and thus for further processing of oxidized nitrogen compounds. However, these reductive pathways do not by themselves explain the formation of NO₂⁻ and NO₃⁻ from NH₄⁺ observed in the ANM assay. Moreover, the canonical denitrification nitrite reductase genes *nirK* and *nirS* were not detected. Therefore, the physiological data and genomic annotations together support a capacity for ammonium utilization and broader inorganic nitrogen transformation in KR4-4ᵀ.

### Nitrogen source utilization characteristics of strain KR4-4ᵀ

To further characterize the nitrogen-utilization phenotype observed during ammonium removal in ANM, the ability of strain KR4-4ᵀ to utilize ammonium, nitrate, and nitrite as individual inorganic nitrogen sources was evaluated using HNM, ADM1, and ADM2, respectively (Figure 4).

**Figure 4.**
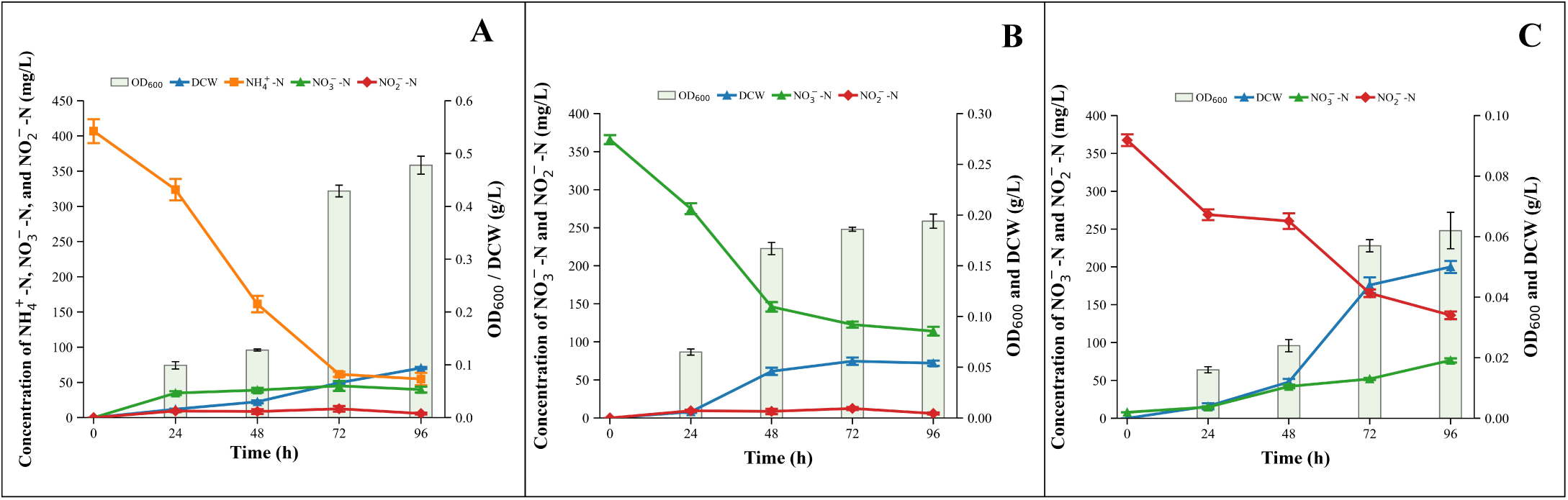
Growth characteristics and nitrogen utilization profiles of strain KR4-4ᵀ with different inorganic nitrogen sources. (A) Ammonium utilization; (B) nitrate utilization; and (C) nitrite utilization.

#### Utilization of ammonium as the sole nitrogen source

When NH₄⁺-N was supplied as the sole nitrogen source in HNM, strain KR4-4ᵀ exhibited progressive growth accompanied by substantial ammonium removal (Figure 4a). The NH₄⁺-N concentration decreased from 406.83 ± 16.99 mg/L at 0 h to 54.84 ± 8.90 mg/L at 96 h, corresponding to a removal efficiency of 86.52% and an average removal rate of 3.67 mg/L/h. This rate was comparable to that reported for *Aeribacillus pallidus* GW-E (3.79 mg/L/h) (30) and higher than those reported for *Bacillus simplex* h-b (0.74 mg/L/h) (31). Concurrently, OD₆₀₀ increased to 0.478 and DCW reached 0.094 g/L, confirming biomass production during ammonium utilization.

Interestingly, NO₃⁻-N and NO₂⁻-N accumulated transiently during cultivation, reaching maximum concentrations of 45.09 ± 1.00 and 12.55 ± 1.97 mg/L, respectively, at 72 h, before decreasing to 39.77 ± 1.15 and 5.88 ± 1.41 mg/L at 96 h. The simultaneous depletion of NH₄⁺-N, increase in biomass, and transient accumulation of NO₃⁻-N and NO₂⁻-N indicate that ammonium removal was associated with both cellular assimilation and transformation into oxidized nitrogen forms. Assimilation into biomass has been reported to account for a substantial fraction of ammonium removal in aerobic heterotrophic nitrogen-removing bacteria (5, 32), whereas the transient formation of nitrate and nitrite observed in KR4-4ᵀ suggests that additional nitrogen-transformation processes may also contribute to its ammonium-removal phenotype.

#### Utilization of nitrate as the sole nitrogen source

When NO₃⁻-N was supplied as the sole nitrogen source in ADM1, strain KR4-4ᵀ exhibited growth accompanied by substantial nitrate removal (Figure 4b). The NO₃⁻-N concentration decreased from 366.67 ± 5.86 mg/L at 0 h to 112.39 ± 5.73 mg/L at 96 h, corresponding to a removal efficiency of 69.35% and an average removal rate of 2.65 mg/L/h. This rate was higher than that reported for *Aeribacillus pallidus* GW-E (1.93 mg/L/h) (30). Concurrently, OD₆₀₀ increased from 0 to 0.194, while DCW increased to a maximum of 0.056 g/L at 72 h and remained at 0.054 g/L at 96 h, demonstrating biomass formation during nitrate utilization.

Nitrite accumulated transiently during nitrate removal. NO₂⁻-N increased from undetectable levels at 0 h to 7.41 ± 1.86 mg/L at 24 h and 8.91 ± 2.40 mg/L at 48 h, reaching a maximum of 32.71 ± 1.64 mg/L at 72 h before decreasing to 19.56 ± 2.00 mg/L at 96 h. The transient accumulation of nitrite is consistent with nitrate reduction to nitrite, an initial step shared by assimilatory and dissimilatory nitrate reduction pathways (5, 8). Its subsequent decline despite the continued presence of nitrate suggests that nitrite underwent further transformation rather than accumulating as a terminal product.

This phenotype is consistent with the genomic repertoire of KR4-4ᵀ, which includes genes associated with nitrate and nitrite transport and reduction, including NRT2, *nirC*, *nasC*, *nasDE*, *narGHI*, and *nrfAH*. Together with the observed biomass formation, these findings support the ability of KR4-4ᵀ to utilize nitrate as a nitrogen source and suggest that nitrate-derived nitrite may subsequently undergo further reduction and/or assimilation.

#### Utilization of nitrite as the sole nitrogen source

Strain KR4-4ᵀ was also able to grow when NO₂⁻-N was supplied as the sole nitrogen source in ADM2 (Figure 4c). The NO₂⁻-N concentration decreased from 367.50 ± 7.74 mg/L at 0 h to 136.18 ± 5.03 mg/L at 96 h, corresponding to a removal efficiency of 62.94% and an average removal rate of 2.41 mg/L/h. This rate was lower than that reported for *Aeribacillus pallidus* GW-E (4.31 mg/L/h) (30) but higher than that reported for *Bacillus* N31 (0.73 mg/L/h) (33). Despite the substantial decrease in NO₂⁻-N, growth was relatively limited, with OD₆₀₀ and DCW reaching only 0.062 and 0.050 g/L, respectively, at 96 h. The limited biomass formation may be associated with the relatively high nitrite concentration, as elevated nitrite levels can impose cytotoxic and oxidative/nitrosative stress on bacterial cells (5).

Notably, NO₃⁻-N increased from 7.97 ± 0.18 mg/L at 0 h to 76.61 ± 2.76 mg/L at 96 h, corresponding to a net increase of 68.64 mg/L. The simultaneous decrease in NO₂⁻-N and accumulation of NO₃⁻-N suggests that a portion of the supplied nitrite was oxidized to nitrate. However, nitrate accumulation accounted for only part of the observed decrease in NO₂⁻-N, indicating that nitrite removal could not be explained solely by its conversion to nitrate. Other processes, including cellular assimilation and additional nitrogen transformations, may therefore have contributed to the overall nitrite-removal phenotype.

The genomic data provide partial support for these observations. Genes associated with nitrite transport and reduction, including *nirC*, *nasDE*, and *nrfAH*, were identified, supporting the potential utilization of nitrite through reductive pathways and subsequent assimilation. However, these genes do not directly account for the observed accumulation of nitrate from nitrite. Thus, while the phenotype demonstrates that KR4-4ᵀ can utilize nitrite as a sole nitrogen source, the enzymatic mechanism underlying the apparent oxidation of nitrite to nitrate remains unresolved based on the current genomic annotation.

Comparison of the three nitrogen-source assays revealed a substrate-dependent response of strain KR4-4ᵀ to inorganic nitrogen. Among the tested substrates, ammonium supported the greatest biomass production and showed the highest removal efficiency (86.52%), followed by nitrate (69.35%) and nitrite (62.94%). The stronger growth response observed with ammonium is physiologically consistent with its direct incorporation into cellular nitrogen metabolism through the GS–GOGAT and GDH pathways. In contrast, nitrate and nitrite must first undergo reduction before their nitrogen can enter these assimilatory routes (5, 32, 8). This interpretation is supported by the genomic repertoire of KR4-4ᵀ, which includes genes for ammonium transport and assimilation (*amt*, *glnA*, *gltBD*, *gdhA*, and *gudB*), together with genes associated with nitrate/nitrite transport and reduction (*NRT2*, *nirC*, *nasC*, *nasDE*, *narGHI*, and *nrfAH*). The particularly limited growth observed with nitrite may also reflect the physiological stress imposed by the relatively high initial nitrite concentration.

Importantly, nitrogen removal in these assays cannot be attributed solely to biomass assimilation. During growth on ammonium, the formation of nitrate and nitrite indicates transformation of a portion of the supplied nitrogen into oxidized forms, whereas nitrate utilization was accompanied by transient nitrite accumulation followed by its subsequent decline. Conversely, nitrite utilization resulted in substantial nitrate accumulation, indicating that part of the supplied nitrite underwent oxidation. These substrate-dependent changes in soluble nitrogen species show that KR4-4ᵀ cannot only assimilate inorganic nitrogen but also transform it among different oxidation states. Nevertheless, the present experiments cannot quantitatively resolve the relative contributions of assimilation and other transformation processes because they did not measure cellular nitrogen and gaseous nitrogen products.

Taken together, the physiological and genomic evidence supports a versatile inorganic nitrogen-utilization phenotype in KR4-4ᵀ, with ammonium supporting the strongest growth under the tested conditions. The genomic potential for assimilatory and dissimilatory nitrate/nitrite reduction is broadly consistent with the observed utilization of oxidized nitrogen sources.

### Gas-phase ammonia reduction by strain KR4-4ᵀ

Given the ability of strain KR4-4ᵀ to remove ammonium and utilize multiple inorganic nitrogen sources in liquid culture, its potential to reduce gaseous ammonia was further evaluated in a closed-flask assay with an initial NH₃ concentration of approximately 1,000 ppm (Figure 5). In flasks inoculated with KR4-4ᵀ, gaseous NH₃ decreased rapidly from 1,000.00 ± 20.52 ppm at 0 h to 423.33 ± 14.53 ppm at 24 h and 280.00 ± 30.55 ppm at 48 h, followed by a more gradual decline to 175.00 ± 21.89 ppm at 96 h. These changes corresponded to apparent reductions in gas-phase NH₃ of approximately 72.0% and 82.5% after 48 and 96 h, respectively. In contrast, NH₃ concentrations in the uninoculated control remained comparatively high, decreasing only from 1,000.00 ± 19.35 to 893.33 ± 12.02 ppm over 96 h. The markedly greater decrease observed in the inoculated treatment indicates that the presence of KR4-4ᵀ substantially enhanced the removal of NH₃ from the gas phase under the tested conditions.

**Figure 5.**
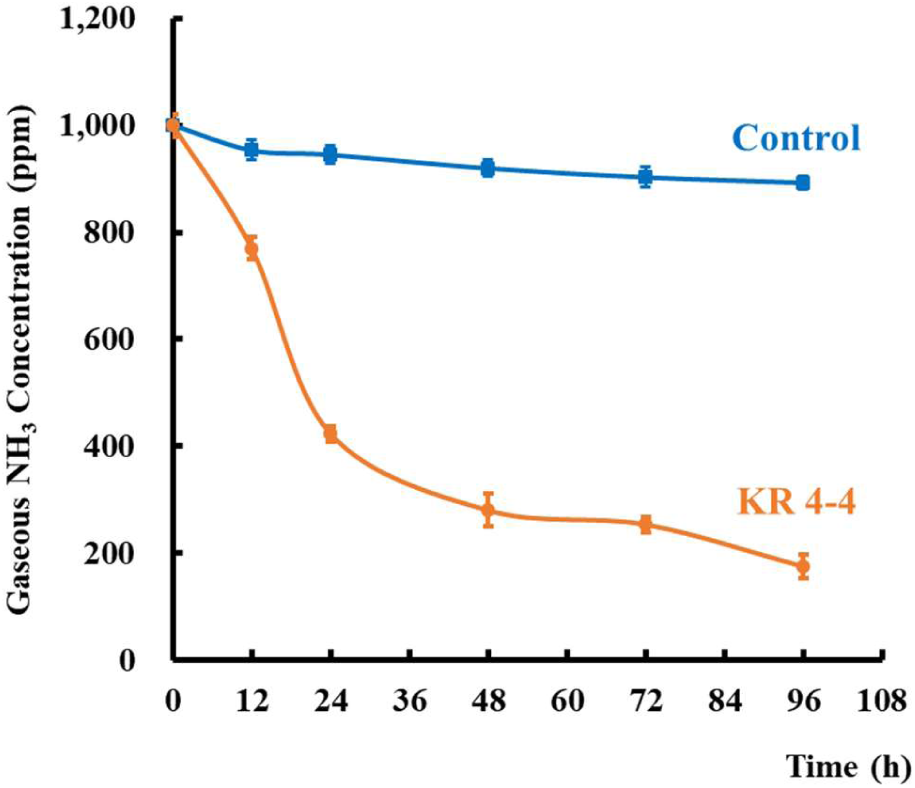
Gas-phase ammonia reduction by strain KR4-4ᵀ in a sealed-flask system.

Biological mitigation of gaseous NH₃ has previously been demonstrated using ammonia-tolerant bacteria and microbial biofiltration systems, highlighting the potential contribution of microorganisms to the control of ammonia emissions and associated malodors (34). In the present study, the ability of KR4-4ᵀ to reduce gas-phase NH₃, together with its capacity to remove NH₄⁺-N and utilize ammonium, nitrate, and nitrite in liquid culture, suggests potential applicability in nitrogen-rich environments where both dissolved and gaseous forms of ammonia may occur. However, the closed-flask experiment does not resolve the fate of the NH₃ removed from the gas phase. Nevertheless, the substantially greater gas-phase NH₃ reduction in the presence of KR4-4ᵀ compared with the uninoculated control supports its further evaluation as a candidate for biological ammonia-mitigation applications.

## Conclusion

Strain KR4-4ᵀ, isolated from soil in the Republic of Korea, represents a novel species of the genus *Neobacillus*, for which the name *Neobacillus ammoniiphilus* sp. nov. is proposed. Its taxonomic status was supported by phylogenetic and phylogenomic analyses, genome relatedness indices, and phenotypic and chemotaxonomic characteristics. Beyond its taxonomic novelty, genomic analysis revealed a repertoire of genes associated with inorganic nitrogen transport, nitrate and nitrite reduction, and ammonium assimilation. Consistent with this genomic potential, KR4-4ᵀ was able to utilize ammonium, nitrate, and nitrite as nitrogen sources and exhibited substantial ammonium removal under the tested conditions. The strain also contributed to the reduction of gaseous NH₃ in a closed-flask assay, further indicating its potential for ammonia mitigation. Collectively, these findings establish KR4-4ᵀ as the type strain of *Neobacillus ammoniiphilus* sp. nov. and expand the known physiological and functional diversity of the genus *Neobacillus*, particularly with respect to inorganic nitrogen utilization and transformation.

### Description of *Neobacillus ammoniiphilus* sp. nov

*Neobacillus ammoniiphilus* (am.mo.ni.i’phi.lus. N.L. neut. n. *ammonium*, ammonium; N.L. masc. adj. *philus*, loving; N.L. masc. adj. *ammoniiphilus*, ammonium-loving, referring to the ability of the type strain to utilize ammonium as a nitrogen source).

Cells are Gram-stain-positive, non-motile, facultatively anaerobic rods, 0.5–1.5 µm wide and 0.9–4.8 µm long. Growth occurs at 15–45 °C (optimum, 30–40 °C), at pH 6.0–8.0 (optimum, pH 7.0-7.5), and in the presence of 0–2.0% (w/v) NaCl. Catalase and oxidase activities are negative. Starch, DNase, aesculin, and gelatin are hydrolysed, whereas casein and urea are not. Nitrate is reduced to nitrite. In API ZYM tests, positive reactions are observed for esterase (C4), esterase lipase (C8), leucine arylamidase, α-chymotrypsin, β-galactosidase, β-glucuronidase, and α-glucosidase. Whereas reactions for lipase (C14), cystine arylamidase, trypsin, β-glucosidase, *N*-acetyl-β-glucosaminidase, α-mannosidase, and α-fucosidase are negative. In API 20NE and API 50CH tests, D-glucose, D-mannose, D-mannitol, N-acetyl-D-glucosamine, D-maltose, malate, phenylacetic acid, D-fructose, aesculin, D-lactose, D-sucrose, D-trehalose, D-melezitose, starch, and glycogen are assimilated, whereas L-arabinose, potassium gluconate, capric acid, adipic acid, and trisodium citrate are not assimilated. The type strain is capable of utilizing ammonium, nitrate, and nitrite as nitrogen sources under the tested conditions. The major cellular fatty acids (>5% of the total) are iso-C15:0 (35.3%), anteiso-C15:0 (33.5%), C16:0 (5.9%), and anteiso-C17:0 (5.1%). The polar lipid profile comprises diphosphatidylglycerol (DPG), phosphatidylglycerol (PG), and phosphatidylethanolamine (PE), together with minor amounts of an unidentified aminolipid and an unidentified phospholipid. The genomic DNA G+C content of the type strain is 38.5%.

The type strain, KR4-4ᵀ (= KACC 24003ᵀ = CCTCC AB 2025077ᵀ), was isolated from soil collected in Baekam-myeon, Cheoin-gu, Yongin, Gyeonggi, Republic of Korea. The GenBank/EMBL/DDBJ accession number for the 16S rRNA gene sequence of the type strain is PP892614, and the whole-genome shotgun sequence has been deposited in DDBJ/ENA/GenBank under the accession number JBHHJU000000000.

## Materials and Methods

### Strain source and culture medium

Strain KR4-4^T^ =KACC 24003 = CCTCC AB 2025077, isolated from a soil sample collected in Baekam-myeon, Cheoin-gu, Yongin, Gyeonggi, Korea (37°07’44.0’N 127°22’01.0’E). A 3 g soil sample was placed at the bottom of a six-well Transwell plate (Corning Inc., Corning, NY, US) and added to 6 ml ammonia nitrogen media (ANM). Subsequently, 100 µl of the prepared soil suspension was transferred into the insert, and cultivation at 28 °C in a shaking incubator at 100 rpm for 1 week. The composition of the ANM medium was adapted from Zhang et al. (2019) (35), which contained (per liter): 5.0 g Glucose, 0.7 g (NH4)2SO4, 1.0 g NaCl, 0.05 g MgSO4 ·7H2O, 0.5 g K2HPO4 ·2H2O, and 0.25 g FeSO4 ·7H2O, with a pH range of 6.79±0.02. After incubation, the suspension was serially diluted, and an aliquot (100 μL) of each dilution was spread onto an ANM agar plate, which was then incubated at 30 °C for 5 days. We picked single colonies and repeatedly streaked them onto ANM plates under aerobic conditions at 30 °C until pure colonies formed. After isolation, we cultured the strain in TSB medium for preservation. The resulting pure cultures were maintained at −80 °C with 20 % (v/v) glycerol until analysis.

## Molecular identification

Genomic DNA from strain KR4-4^T^ were extracted using InstaGene Matrix kits (Bio-Rad; Hercules, CA, USA) according to the manufacturer’s instructions. The 16S rRNA gene sequences of the strain were amplified using the universal bacterial primer sets 518F/805R and 27F/1492R, and the PCR products were commercially sequenced (Macrogen, Seoul, Republic of Korea). The obtained 16S rRNA gene sequence was submitted to the EzBioCloud server (https://www.ezbiocloud.net/) (36) for species identification by comparison with closely related bacterial type strains. The 16S rRNA gene sequences of the closest phylogenetic relatives were aligned using SINA (https://www.arb-silva.de/aligner/) (37). We conducted phylogenetic analyses using 16S rRNA gene sequences from the novel strains and related references in MEGA 11 (38). Phylogenetic trees were reconstructed using the maximum likelihood (ML), minimum evolution (ME), and neighbour-joining (NJ) methods, applying Kimura’s two-parameter method with 1,000 bootstrap replications. The sequences of Bacillus subtilis 168T (NR102783) were used as outgroups.

## Phenotypic and chemotaxonomic characterization

The cell morphology and size of strain KR4-4^T^ was observed using transmission electron microscopy (Talos L120C, FEI). Gram staining was performed using a commercial reagent kit (Sigma, St. Louis, MO, United States). Catalase oxidase test were assessed using 3% (v/v) hydrogen peroxide and 1 % (w/v) tetramethyl-p-phenylenediamine (bioMérieux), respectively. The temperature tolerance and the optimum growth temperature were tested on R2A plates at various temperatures, ranging from 4 to 55 °C (4, 10, 15, 20, 25, 30, 35, 40, 45, 50 and 55 °C) over 5 days. Salt tolerance and the optimal NaCl concentration for growth were evaluated on R2A broth containing NaCl concentrations ranging from 0 to 10.0% (at 0.5% increments) for 5 days at 30 °C. The pH range for growth and the optimal pH value were determined using media with pH ranging from 4.0 to 11.0 at 0.5 pH unit increments, following the buffer system as previously described (39). The hydrolysis of starch (Duksan), chitin (Tokyo Chemical Industry Co., Ltd.), casein (1 % skimmed milk; MB cell), Tween 80 (1 %; Sigma-Aldrich) were tested on R2A agar medium for 5 days at 30 °C. Other biochemical and physiological tests were determined using API ZYM, API20 and API50CH test kits (bioMérieux) according to the manufacturer’s instructions.

For chemotaxonomic analysis, strain KR4-4^T^ was grown on TSB agar for 3 days at 30°C until reaching the post-growth stage. Polar lipids were extracted from freeze-dried cells according to the procedures described by Minnikin et al. (1984) (40). Lipids were separated by two-dimensional chromatography on a silica gel thin-layer chromatography plate (Kieselgel 60 F254; Merck). Cellular fatty acids analysis was extracted using MIDI protocol (41) and analysed using a gas chromatograph (GC; HP 6890 Series GC System; Hewlett Packard) equipped with Sherlock Microbial Identification System software V6.01 (MIS, database TSBA6, midi Inc.)

### Genome sequencing, assembly, annotation and analysis

Whole-genome sequencing of KR4-4^T^ was performed by Macrogen Co., Ltd. (Seoul, Republic of Korea). Genomic DNA was extracted from 48-h pure cultures grown on TSB medium using the DNeasy Blood and Tissue Kit (Qiagen). Sequencing libraries were prepared with the TruSeq Nano DNA High Throughput Library Prep Kit (Illumina) following the manufacturer’s protocol. Library concentration was determined using KAPA Library Quantification kits for Illumina sequencing platforms (KAPA BIOSYSTEMS, #KK4854) according to the qPCR Quantification Protocol Guide and library quality was assessed using the TapeStation D1000 ScreenTape (Agilent Technologies, # 5067-5582). The indexed libraries were subsequently sequenced on an Illumina MiSeq platform (Illumina, Inc., San Diego, CA, USA) to generate paired-end reads (2 × 301 bp). Raw sequencing reads were error-corrected using the k-mer-based algorithm implemented in the A5-miseq pipeline (v.20160825), and adapter sequences were removed with Trimmomatic v0.38. Read quality was evaluated using FastQC 0.11.7 (42). De novo assembly was performed with SPAdes 3.15.0 (43) using multiple k-mer sizes. Contigs shorter than 1,000 bp were discarded to minimize assembly bias, and the completeness of the assembled genome was assessed using BUSCO v5.1.3 (44) against the bacterial ortholog database (ODB10).

Phylogenomic analysis of strain KR4-4T and related members of the genus *Neobacillus* was performed using the UBCG pipeline (45), based on the concatenated alignment of 92 conserved bacterial core genes. Core genes were identified with the default UBCG parameters, aligned using MAFFT, and concatenated for phylogenomic tree reconstruction with RAxML. The resulting tree was visualized using MEGA 11 (38). The whole-genome sequence of *Bacillus subtilis* 168^T^ were used as the outgroup. To analyse genome-based relatedness between strain KR4-4^T^ and the related strains, genome sequences of the reference strains were collected from the NCBI database used for comparisons with the novel strains for average nucleotide identity (ANI) using the Orthologous Average Nucleotide Identity Tool from EzBioCloud (46) and digital DNA–DNA hybridization (dDDH) values were determined using the Type (Strain) Genome Server (https://tygs.dsmz.de) (47).

Genome annotation was performed using the NCBI Prokaryotic Genome Annotation Pipeline (PGAP). Functional annotation of the predicted protein-coding genes was further conducted using the RAST annotation server (48), BlastKOALA (49) and antiSMASH server version 7.0 (50). Metabolic pathways were reconstructed using KEGG Mapper (51) based on the KO assignments generated by BlastKOALA.

### Nitrogen utilization assays

The nitrogen utilization characteristics of strain KR4-4ᵀ were evaluated by examining ammonium removal and the utilization of different inorganic nitrogen sources. Unless otherwise specified, cultures were incubated at 35 °C and 180 rpm for 96 h, uninoculated media incubated under the same conditions were used as abiotic controls, and all experiments were performed in triplicate.

#### Ammonium removal

The ammonium removal capacity of strain KR4-4ᵀ was evaluated using ammonium nitrogen medium (ANM). For inoculum preparation, strain KR4-4ᵀ was cultured in ANM broth at 35 °C and 180 rpm for 48 h. Subsequently, 6 mL of the seed culture was inoculated into 300 mL of ANM containing an initial NH₄⁺-N concentration of 184 ± 3 mg L⁻¹ in a 500-mL Erlenmeyer flask. Samples were collected at 0, 12, 24, 36, 48, 60, 72, 84, and 96 h to determine cell growth (OD₆₀₀) and the concentrations of NH₄⁺-N, NO₃⁻-N, and NO₂⁻-N.

#### Utilization of different inorganic nitrogen sources

The ability of strain KR4-4ᵀ to utilize different inorganic nitrogen sources was evaluated using heterotrophic nitrification medium (HNM) and aerobic denitrification media (ADM1 and ADM2), prepared according to Zhang et al. (2025) (30). HNM contained ammonium as the sole nitrogen source, whereas ADM1 and ADM2 contained nitrate and nitrite as the sole nitrogen sources, respectively. The compositions of the media and their initial nitrogen concentrations are provided in Table S8.

For inoculum preparation, strain KR4-4ᵀ was cultured in tryptic soy broth (TSB) at 35 °C and 180 rpm for 48 h. Cells were harvested by centrifugation at 15,000 rpm for 5 min at 4 °C, washed twice with sterile distilled water, and resuspended in sterile distilled water. The resulting cell suspension was inoculated at 1% (v/v) into Erlenmeyer flasks containing 200 mL of HNM, ADM1, or ADM2. Samples were collected at 0, 24, 48, 72, and 96 h to determine cell growth (OD₆₀₀), dry cell weight (DCW), and the concentrations of NH₄⁺-N, NO₃⁻-N, and NO₂⁻-N.

#### Analytical methods

Cell growth was monitored by measuring the optical density at 600 nm (OD₆₀₀). NH₄⁺-N was determined using the Nessler reagent colorimetric method, whereas NO₃⁻-N and NO₂⁻-N were determined by ion chromatography (Dionex™ ICS-1600 equipped with an AS14 column). Dry cell weight (DCW) was determined by centrifugation, followed by washing the cell pellets with distilled water and drying at 80 °C until a constant weight was obtained (52).

## Calculation of nitrogen removal

Nitrogen removal efficiency and average removal rate were calculated from the initial and final concentrations of the respective nitrogen compounds according to Zhang et al. (30), as follows:

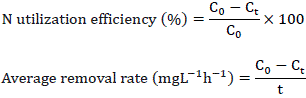

where C0 and Ct represent the concentrations (mg L⁻¹) of the respective nitrogen compound at the beginning of incubation and after t hours, respectively.

## Gaseous ammonia reduction assay

The ability of strain KR4-4ᵀ to reduce gaseous ammonia (NH₃) was evaluated according to Bajagain et al. (2022) (38), with minor modifications. The assay was conducted in 250-mL Erlenmeyer flasks containing tissue paper moistened with an ammonia solution to generate gaseous NH₃. The initial NH₃ concentration inside each flask was adjusted to 1,000 ± 50 ppm. Subsequently, 10 mL of a 48-h bacterial culture grown in TSB was sprayed into each flask. The flasks were immediately sealed and incubated at 30 °C without agitation. For the control, 10 mL of sterile TSB was applied to otherwise identical flasks and incubated under the same conditions. Gaseous NH₃ concentrations were measured using a Gastec GV-100S sampling pump (Gastec Corporation, Japan) equipped with appropriate NH₃ detector tubes after 12, 24, 48, 72, and 96 h of incubation. All treatments were performed in triplicate.

## Data analysis

Experimental data are presented as the mean ± standard deviation of three replicates. Microsoft Excel was used for data organization and preliminary calculations. Data visualization and figure preparation were performed using Python version 3.13.15.

## Data availability

The type strain is KR4-4^T^ =KACC 24003^T^ = CCTCC AB 2025077^T^. The 16S rRNA gene sequence of strain KR4-4^T^ has been deposited in the NCBI GenBank database under accession number PP892614. The whole-genome shotgun project has been deposited in DDBJ/ENA/GenBank under accession number JBHHJU000000000. The raw sequencing reads have been deposited in the NCBI Sequence Read Archive (SRA) under accession number SRR39355900.

